# A *de novo* long-read genome assembly of the sacred datura plant (*Datura wrightii*) reveals a role of tandem gene duplications in the evolution of herbivore-defense response

**DOI:** 10.1101/2023.05.08.539846

**Authors:** Jay K Goldberg, Aaron Olcerst, Michael McKibben, J Daniel Hare, Michael S Barker, Judith Bronstein

## Abstract

The sacred datura plant (Solanales: Solanaceae: *Datura wrightii*) has been used to study plant-herbivore interactions for decades. The wealth of information that has resulted leads it to have potential as a model system for studying the ecological and evolutionary genomics of these interactions. We present a *de novo Datura wrightii* genome assembled using PacBio HiFi long-reads. Our assembly is highly complete and contiguous (N50 = 179Mb, BUSCO Complete = 97.6%). We successfully detected a previously documented ancient whole genome duplication using our assembly and have classified the gene duplication history that generated its coding sequence content. We use it as the basis for a genome-guided differential expression analysis to identify the induced responses of this plant to one of its specialized herbivores (Coleoptera: Chrysomelidae: *Lema daturaphila*). We find over 3000 differentially expressed genes associated with herbivory and that elevated expression levels of over 200 genes last for several days. We also combined our analyses to determine the role that different gene duplication categories have played in the evolution of *Datura*-herbivore interactions. We find that tandem duplications have expanded multiple functional groups of herbivore responsive genes with defensive functions, including UGT-glycosyltranserases, oxidoreductase enzymes, and peptidase inhibitors. Overall, our results expand our knowledge of herbivore-induced plant transcriptional responses and the evolutionary history of the underlying herbivore-response genes.

## Introduction

*Datura wrightii* (Solanales: Solanaceae) has begun to serve as a model system for research into the ecology and evolution of various plant traits, including both physical and chemical defenses [1,2], tolerance to herbivory [3], floral phenotypes [4,5], life histories [6,7], and ontogenetic changes in defense production throughout an individual plant’s lifetime [8,9]. This is due to the wealth of ecological knowledge already gathered regarding this common plant and the specialist insects with which it interacts [10,11]. Furthermore, field-based studies have already provided some insights into the evolutionary processes maintaining a trichome dimorphism in naturally occurring populations [2,12]. As such, the publication of a reference genome for this species would accelerate research into both the molecular mechanisms governing the expression of plant traits and their evolutionary trajectories in a rapidly changing environment.

Here, we present a *de novo* genome assembly for *D. wrightii*, generated using highly accurate long-reads (PacBio HiFi) [13]. Our assembly is highly complete and contiguous when compared to recent assemblies of closely related species [14,15,16]. To demonstrate the utility of our genome to future research, we conducted two downstream analyses with it: 1) an assessment of the duplication history of coding regions with our assembly and 2) a differential expression study examining herbivore-induced transcriptional changes. We further use the results of these analyses to determine the role that different categories of gene duplications have had on the herbivore-induced responses of this species.

Gene and whole genome duplications (WGDs) have broad implications for evolutionary processes including the generation of novel traits [17,18]. WGDs have occurred numerous times throughout the diversification of angiosperms [19]; and these ‘ancient’ WGDs have been shown to underlie the arms-race dynamic occurring between plants and herbivores [20]. There is widespread evidence of an ancient hexaploidy event that occurred early in the evolution of the Solanaceae [19]. As such we used two different analyses to infer and characterize this ancient WGD in our genome to both detect its presence and assess the ancestral ploidy level of the lineage.

We further use our assembly to conduct a differential expression study into the induced responses of *D. wrightii* to one of its specialist herbivores, the three-lined potato beetle (*Lema daturaphila*) [10,11]. This analysis utilized older data, generated roughly a decade ago [21]. Using both pairwise and time course analyses, we identified thousands of genes that are significantly differentially expressed when plants are under attack by one of their closest natural enemies, and further show that many of them remain upregulated for multiple days. We conclude by using an over-representation analysis approach to determine the molecular functions that various gene duplication categories have expanded.

## Methods

### Initial sample collection and preparation, and sequencing

*Datura wrightii* seeds were originally collected in June 2016 from a wild plant growing at the intersection of Portal Rd. and Foothills Rd. in Portal, Arizona, USA (see https://www.inaturalist.org/observations/156947070 for more details). A cohort of seeds from this collection was germinated in May 2021. Leaf tissue collected for genome sequencing originated from a single individual. Tissue samples were flash frozen and ground under liquid nitrogen immediately before storage at -80C. Samples for differential expression analysis were collected from two experiments conducted at UC Riverside in 2012/13 [21]. Both studies were conducted with the MVV6 line, a backcrossed line originating from seeds originally collected from Moreno Valley, CA that induces production of defenses in response to herbivore attack [36].

### Nucleic acid extractions and sequencing strategies

DNA extraction, SMRT bell library preparation, and sequencing (PacBio HiFi, Pacific Bioscience, San Francisco, CA, USA) were performed by the Arizona Genomics Institute (University of Arizona, Tucson, AZ, USA). High molecular weight DNA was extracted from young leaves using the protocol of Doyle and Doyle [22] with minor modifications. Flash-frozen young leaves were ground to a fine powder in a frozen mortar with liquid nitrogen followed by very gentle extraction in 2% CTAB buffer (that included proteinase K, PVP-40 and beta-mercaptoethanol) for 30min to 1h at 50 °C. After centrifugation, the supernatant was gently extracted twice with 24:1 chloroform:isoamyl alcohol. The upper (aqueous) phase was then removed and 1/10^th^ volume 3 M NaAc was added, gently mixed, and then had DNA precipitated with iso-propanol. DNA was collected by centrifugation, washed with 70% ethanol, air dried for 20 minutes and dissolved thoroughly in elution buffer at room temperature followed by RNAse treatment. DNA purity was measured with Nanodrop, DNA concentration measured with Qubit HS kit (Invitrogen) and DNA size was validated by Femto Pulse System (Agilent).

DNA was sheared to an appropriate size range (10-20 kb) using Megaruptor 3 (Diagenode) followed by Ampure bead purification. The sequencing library was constructed following manufacturers protocols using SMRTbell Prep kit 3.0. The final library was size selected on a Pippin HT (Sage Science) using S1 marker with a 10-25 kb size selection. The recovered final library was quantified with Qubit HS kit (Invitrogen) and size checked on Femto Pulse System (Agilent). The final library was prepared for sequencing with PacBio Sequel II Sequencing kit 2.0 for HiFi library, loaded on 8M SMRT cells, and sequenced in CCS mode for 30 hours.

RNA was extracted from root and bud tissue using a ZYMO (Irvine, CA, USA) direct-zol miniprep kit (Cat. # R2050) and sequenced using NovaSeq (Illumina, San Diego, CA, USA) paired-end (150bp) sequencing performed by Novogene (Sacramento, CA, USA). RNA for differential expression was extracted and sequenced as described in Olcerst (2017) [21].

### Genome assembly/annotation

CCS output (ie: HiFi reads; 3952061 reads; 65.26Gb total; mean length = 16524) were assembled using hifiasm-0.16.0 [23] with default settings. Jellyfish v2.2.10 [31] was used for kmer counting (kmer size = 101bp) before using the GenomeScope2.0 web portal [41] to estimate genome size (Figure S1). Genome quality was examined using Bandage v0.8.1 [27], BUSCO v5.1.3 (odb10_solanales; Figure 1) [28], the blobtools v1.1 pipeline (Figure S2) [29] employing minimap v2-2.24 [30] for alignment and the nt sequence database for taxonomic identification (BLAST 2.13.0) [42], and Inspector [43]. Inspector was also used for error correction/polishing (Table S1). Repetitive elements were identified using RepeatModeler v2.0.1 and RepeatMasker v4.1.0 [24] (Table S2). Structural gene annotation was carried out using the Helixer v0.3.1 algorithm pipeline [44,45] using the pre-made land plant training dataset. Functional annotation was done using InterProScan v5.45-80.0 [46] and blastp (using blast v2.13.0) [42] comparisons to the UniProt-Swissprot database [47]. Annotation files were combined into a single gff using the manage_functional_annotation.pl script in the AGAT v 1.2.0 toolkit [48]. Detailed annotation statistics are shown in Table S3.

**Figure 1.**
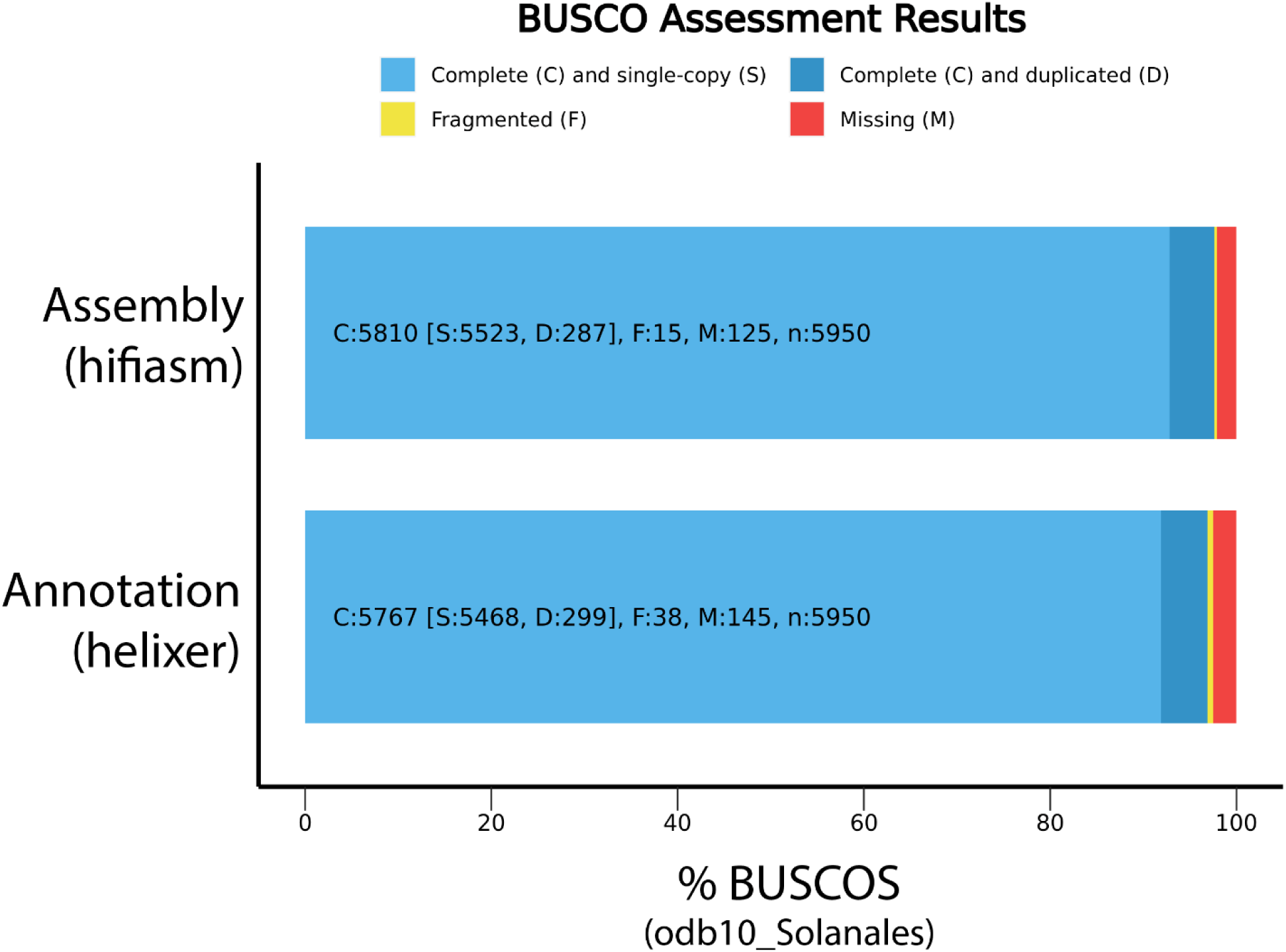
Bar charts showing the gene content represented in both our assembly (top bar) and our annotation (bottom bar). Annotation assessment was run in proteome mode.

### RNA-seq read alignments

Raw RNA-seq reads were aligned to the annotated reference genome and counted using STAR [37] with the default parameters. STAR was selected for its flexible alignment parameters (up to 10 mismatches per read, so long as total mismatches do not exceed 30% of their length), and automatic read counting functionality when given an annotation (a .gff3 file in our case). Reads aligned to our reference genome at an average rate of 80% per library, with a multi-mapping rate of less than 10% for all libraries. An average of 0.3% of reads from each library was removed for mapping to over 10 loci. Alignment details can be found in Table S4. Untransformed read counts were passed to DESeq2 [38] in R [39] for statistical analysis of differential gene expression.

### Gene duplication categorization

We used two programs, MCScanX [32] and frackify [49], to investigate the history of gene duplication events that generated our observed *D. wrightii* gene content and test for signatures of ancient WGDs. Using MCScanX we made inter- and intraspecific syntenic comparisons of the *D. wrightii* and *Ipomoea purpurea* reference genomes [32]. We visualized the syntenic depth ratio of each collinear loci in the interspecific comparison using matplotlib [35]. Additionally we appended Ka/Ks values for each collinear gene pain in both the inter- and intraspecific syntenic comparison using the add_kaks_to_synteny.pl script available through MCScanX [32]. The distribution of Ks values for each comparison were transformed into density functions with the nparam_density and gaussian_kde functions from the Numpy and Scipy python libraries [59]. Maxima of each WGD and ortholog divergence peak in each density distribution were found with the find_peaks function from the Scipy python library [59]. We then used the syntenic and Ks inferences as input for Frackify to identify paleologs in the D. wrightii reference genome [49]. Finally, we classified other paralogs in the genome as tandem, dispersed, proximal, and segmental duplicates using duplicate_gene_classifier available through MCScanX [32].

### Differential expression experiment 1: pairwise comparison & gene set enrichment analysis

Adults and larvae of *L. daturaphila both* feed on the leaves of *D. wrightii.* Both life stages remove leaf tissue in irregular holes between the veins (Hare & Elle 2002). The overall lifecycle of the beetle is about a month, and the duration of its larval period is about a week, depending upon temperature. Many generations of *L. daturaphila* are produced over the nine-month growing season of *D. wrightii.* Our pairwise experiment consisted of three replicates of control/ *L. daturaphila*-induced leaf samples (N = 6 total RNAseq libraries) collected from greenhouse grown plants in May 2012. Samples were collected after *L. daturaphila* larvae had been allowed to feed for 24h. Additional details regarding growing conditions and sample collection can be found in Olcerst [21]. We used the Wald test in DESeq2 [38] to test for pairwise differences between control and *L. daturaphila*-challenged samples while accounting for individual differences between sample pairs. We used a significance cutoff of P_adj_=0.05 for all gene-wise analyses without any fold-change cutoff for differential expression. We then performed a gene set enrichment analysis (GSEA) on the results of our pairwise experiment using the ClusterProfiler 4.0 package [50]. Each GO ontology (biological processes, molecular functions, and cellular component) was analyzed separately. We further separated each ontology into separate up- and down-regulated gene lists as prior studies have found this approach to be more robust than grouping all DEGs [51].

### Differential expression experiment 2: time course

The quantities and composition of volatile compounds induced in *D. wrightii* by *L. daturaphila* both vary with the time after herbivore damage [57]. This time course study asked how the pattern of gene induction might also vary over time since induction. Our experiment consisted of samples taken at 5 timepoints during larval herbivory (0h [before treatment], 12h, 24h, 48h, 96h; 6 per timepoint with a balanced design; except 24h where N = 5, 3 control, 2 *L. daturaphila*-induced). We used a likelihood ratio test (LRT) approach to analyze these data. Our full model consisted of timepoint, treatment type, and the interaction term. The reduced model lacked the interaction term; thus, this analysis tested the significance of the interaction term specifically (i.e. genes for which expression over time differed by treatment). We then used the DEGreport package [52] to identify co-expressed gene groups within the time course dataset and visualize their expression levels over time.

### Duplicated gene over-representation analyses

To determine the role of gene duplication categories in expanding functional gene groups, we used an over-representation analysis (ORA) approach via the ClusterProfiler package in R [50]. This tests for enriched GO terms within a subset of genes compared to a background set of genes. We only used the molecular function ontology for this analysis. We analyzed each gene duplication category as separate subsets against the total set of genes with assigned GO terms (N = 24939). We also conducted this analysis using only the set of significantly differentially expressed genes with assigned GO-terms (N = 1954).

## Results

### Genome assembly and annotation

Our *de novo* assembly is highly complete and contiguous (N50 = 179Mb; Longest contig = 202Mb; 1144 contigs; BUSCO odb10_Solanales Complete=97.7% [Single-copy=92.9%, Duplicated=4.8%], Fragmented=0.2%, Missing=2.1%; Figure 1; Table 1). The total length of our assembly is 2.2Gb, larger than other *Datura* assemblies [14,15] and close to the prediction obtained from GenomeScope (2.086 Gb; Figure S1). When compared to previous *Datura* assemblies, we find that our assembly contains far fewer contigs that are orders of magnitude longer (see Table 1). This improvement is due entirely to our use of PacBio HiFi reads, rather than previous generation sequencing technologies. Inspector analysis found several structural errors (N=50) and small-scale errors (N = 37211; 16.63 per Mb) in our assembly. Polishing our assembly did not substantially reduce the number of structural errors (N_polished_ = 49), but did reduce the number of small-scale errors in our assembly (N_polished_ = 2808; 1.255 per Mb). Detailed output from inspector, before and after polishing, are found in Table S1. Blobtools analysis determined that no contamination was present in our assembly and that all contigs/reads were identified as belonging to the Solanaceae (Figure S2). 86.11% of the assembly is repetitive elements, primarily retroelements such as long terminal repeats (Table S2). Structural gene annotation using the helixer pipeline was able to preserve the majority of gene content represented in our assembly (BUSCO odb10_Solanales: Complete=96.9%, [Single-copy=91.9%, Duplicated=5.0%], Fragmented=0.6%, Missing=2.5.8%; N_genes_ = 45500; Figure 1). Of the structurally annotated genes, 42745 were functionally annotated; 37040 of which have inferred gene names from blastp against the Uniprot/Swissprot database [47]. Detailed annotation statistics are found in Table S3.

**Table 1.** Summary of assembly statistics compared to those of *Datura stramonium* genomes previously published by De-La-Cruz et al. (2021) [15].

|  | *D. wrightii*<br>(Portal, AZ,<br>USA) | *D. stramonium*<br>(Ticumán, MX) | *D. stramonium*<br>(Teotihuacán, MX) |
| --- | --- | --- | --- |
| **Total Size (Gb)** | 2.2 | 1.48 | 1.28 |
| **Contig number** | 1144 | 27915 | 30392 |
| **Largest contig (Mb)** | 202 | 3.13 | 2.11 |
| **N50** | 179Mb | 84.1kb | 58.2kb |
| **N75** | 121Mb | 44.5kb | 32.6kb |
| **L50** | 5 | 4557 | 5713 |
| **L75** | 10 | 10641 | 13166 |
| **BUSCO complete (%)** | 97.70 | 91 | 81.70 |
| **BUSCO db (odb10)** | Solanales | Solanaceae | Solanaceae |

### Gene Duplication Analyses

We used MCScanX [32] to test for the presence of the ancient Solanaceae WGD [16,19] using a Ks analysis of the colinear gene blocks within our assembly. This analysis identified a large peak (Ks = 0.70), indicative of a rapid burst of gene duplications and consistent with the predictions of ancient WGD (Figure 2A). We used a neighboring lineage of the Solanaceae that does not share the ancient WGD (Solanales: Convolvulaceae: *Ipomoea purpurea*) to calculate the syntenic depth of loci collineary to the *D. wrightii* genome assembly (Figure 2B). A significant number of collinear loci (N = 972) had a syntenic depth ratio of 1:3, consistent with paleohexaploid ancestry. These inferences were further validated using Frackify to identify multi-copy paleologs retained from the Ks peak at 0.70. Among the paleologs retained in duplicate, 17% were retained in the triple copy state. We further used MCScanX to classify all the remaining paralogs within the largest 50 contigs of the genome assembly (Table 2; Table S8). We found that dispersed duplications are most common (N = 20523; Table 2), 75% of which were identified as paleologs by Frackify. Ancient WGDs are likely to have generated a significant amount of the gene content observed in the *D. wrightii* genome as well (N = 30714). All of these results are consistent with prior analyses that the ancient Solanaceae WGD was an ancient hexaploidization [16,19].

**Figure 2.**
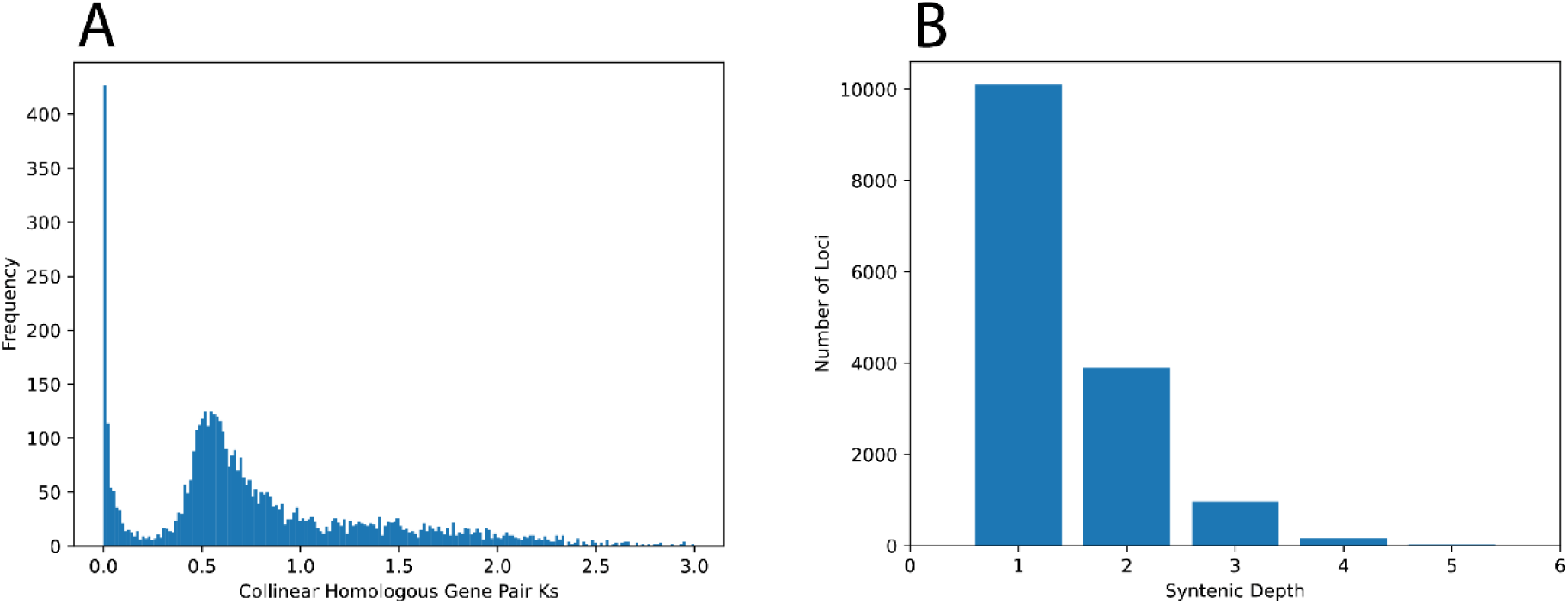
(A) Results of MCSxanX showing the frequency of non-synonymous substitutions in homologous syntenic gene blocks. The large peak at ∼0.5Ks indicates a large burst of gene duplications, consistent with the presence of an ancient whole-genome duplication. (B) Frackify showing the syntenic depth of gene blocks in the *Datura wrightii* genome compared to *Ipopmoea purpurea*. The elevated number of triplicated syntenic blocks indicates an ancient hexaploidy state in *D. wrightii* that is not present in *I. purpurea*.

**Table 2.** Results of MCScanX gene duplication classifier function and GO-term over-representation analysis by duplication type. Gene counts refer to the total number of genes (ie: with or without assigned GO-terms).

| Duplication type | Gene count |  | Enriched GO-terms<br>(Molecular function) |  |
| --- | --- | --- | --- | --- |
|  | Whole<br>genome | DEGs only | Whole<br>Genome | DEGs only |
| **Singleton** | 4268 | 431 | NA | NA |
| **Dispersed** | 20523 | 1329 | 10 | 1 |
| **Proximal** | 2140 | 154 | 28 | 2 |
| **Tandem** | 4064 | 487 | 45 | 8 |
| **Segmental/WGD** | 14505 | 1196 | 13 | 1 |

### Differential expression experiment 1: pairwise comparison & gene set enrichment analysis

Our pairwise analysis found that control and *L. daturaphila*-induced plants had distinct gene expression profiles (Figure 3A) driven by 3555 significantly differentially expression genes (DEGs; Figure 3B; Table S5). Most DEGs (N = 1985) were up regulated in response to *L. daturaphila* feeding (Figure 3B). Up-regulated genes had a far greater range of fold changes (Log_2_FC max = 10.3) than down-regulated genes (Log_2_FC min = -7.3). Functional annotation identified many genes as being involved with known herbivore-response processes such as jasmonic acid signaling and terpene synthesis (Table S5). Gene set enrichment analysis (GSEA) further confirmed this and demonstrated that several functional groups are differentially expressed in *L. daturaphila* -induced plants (Figure 4; Table S6). Terpene synthase activity was notably enriched within our set of up-regulated DEGs alongside other known herbivore-response functions such as peptidase inhibitors, oxidoreductase enzymes, and UDP-glycosyltransferases (Figure 4C). UDP-glycosyltransferase and oxidoreductase activity were also found to be significantly enriched in our set of down-regulated DEGs (Figure 4D). Extracellular and apoplastic gene products were found to be enriched in up-regulated DEGs (Figure 4E), consistent with previous findings in other plant-insect interactions [53]. Gene-wise results of pairwise DEG analysis are found in Table S5 whereas the full GSEA results and summary statistics are shown in Table S6.

**Figure 3.**
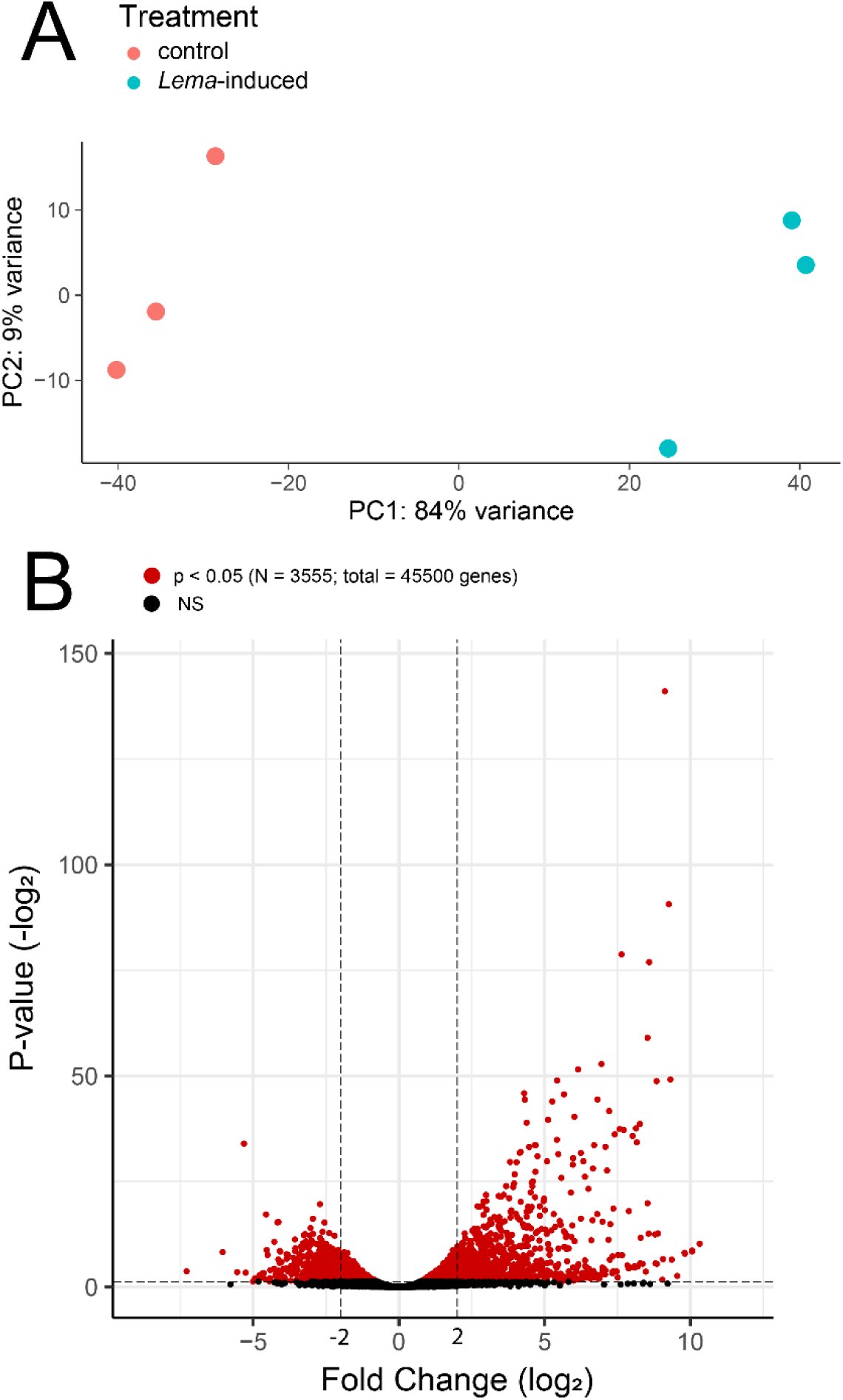
Results of pairwise differential expression analysis using only the samples collected in May 2012. (A) Results of exploratory PCA to qualitatively screen for differences between treatment groups. There was a clear qualitative difference between the control and *L. daturaphila*-induced samples driven by PC1. (B) Volcano plot showing the relationship between significance and fold-change for each gene in our *D. wrightii* genome annotation. A change of 2/-2 is marked for reference but was not used as a cutoff for any analysis. Significant genes (p < 0.05) are shown in red, non-significant genes in black.

**Figure 4.**
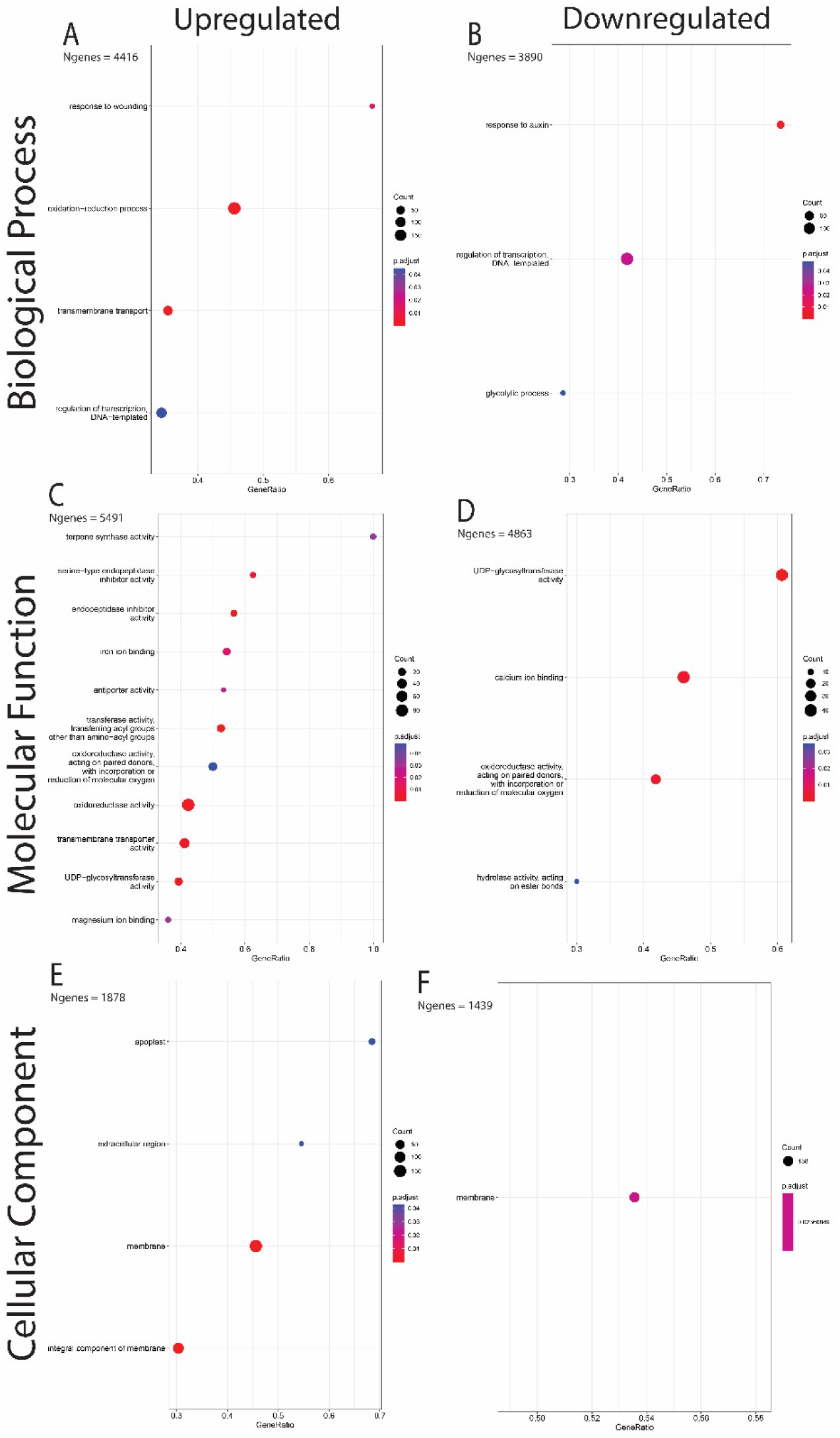
Results of gene set enrichment analysis of pairwise DEGs. X-axes show the ratio of enriched genes vs the total count of genes sharing that GO term. Dot sizes represent the total number of genes sharing each GO term whereas dot colors represent the p-value (adjusted for multiple tests) of each term. Each GO ontology was analyzed separately for up- and down-regulated genes.

### Differential expression experiment 2: time course

Our time course analysis generated a shorter list of differentially expressed genes (N = 605; Figure 5; Table S7). Principle component analysis (PCA) indicated that *L. daturaphila*-induced samples at 24h, 48h, and 96h formed a distinct cluster (Figure 5A). We also noticed a strong differentiation between 0h (pre-herbivory) and 12h post-herbivory treatments (Figure 5A), regardless of treatment type (control or *L. daturaphila*-induced), which may be a sign of background circadian rhythms in gene expression [40]. Most of these genes (N_total_ = 495) clustered into nine different groups based on their expression profiles over time (Figure S3), but only the two largest clusters showed a clear difference between control and *L. daturaphila*-induced treatments (Figure 5B/C). Genes in these clusters were up-regulated in the 24h, 48h, and 96h timepoints, consistent with our PCA results.

**Figure 5.**
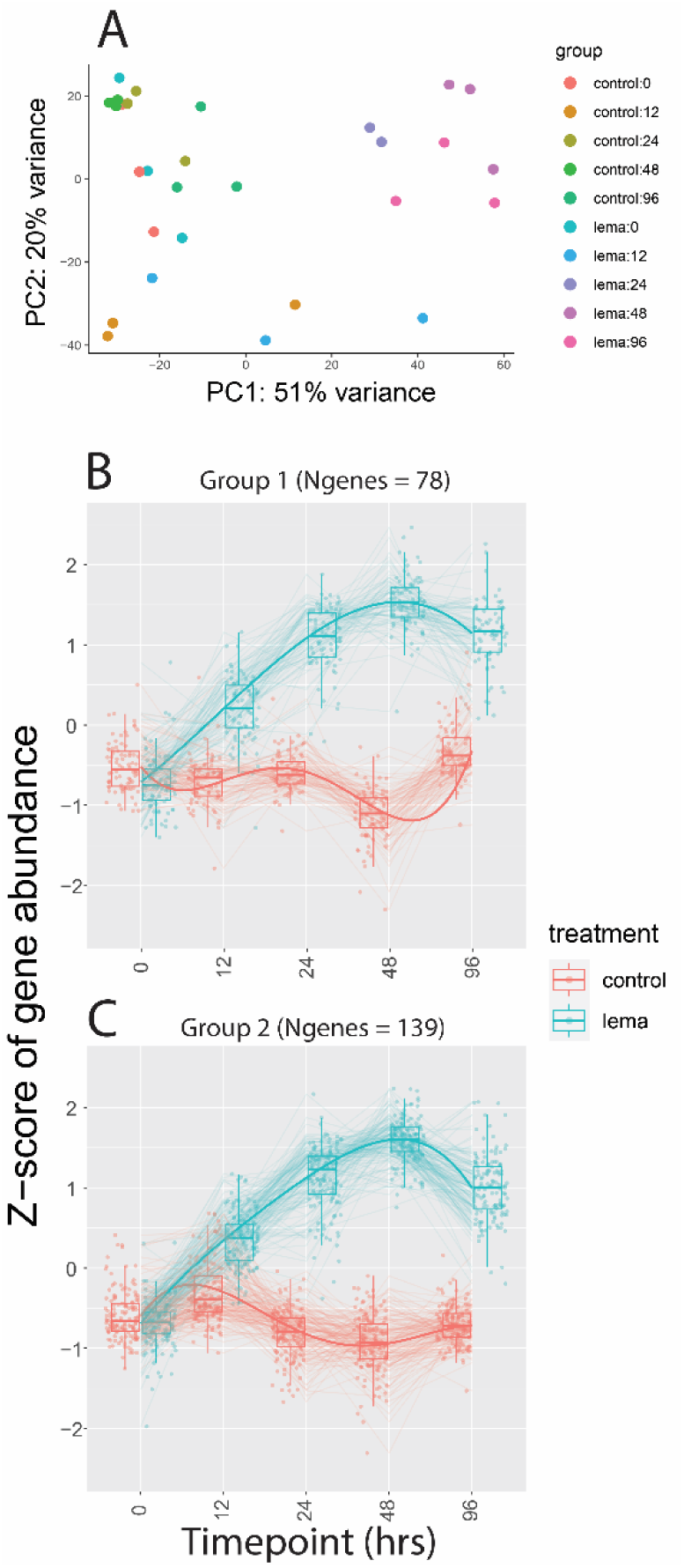
(A) Exploratory PCA of time course data. Although our samples did not form clear clusters, there are definitive patterns. The 24h, 48h, and 96h *L. daturaphila*-induced samples cluster in the top right, whereas the controls for these time points cluster with the 0h samples in the top left. Ranges for 0h and 12h samples do not overlap at all, suggesting that there may also be circadian cycles occurring. (B/C) Cluster plots showing the two largest gene clusters with similar expression patterns. These are the only gene clusters that showed a clear separation between treatments throughout the experiment time. Extended gene cluster results are shown in Figure S3.

### Gene duplication over-representation analysis

When assessing the functional groups enriched by different duplications, we found that tandem duplications have expanded the most molecular functions (N = 45; Table 2; Table S9). Many of these functions – such as chitin binding, hydrolase enzymes, peptidase inhibitors, and terpene synthases – have known roles in defense against herbivores [53,54]. When we analyzed only the enriched functions within our list of DEGs, we again found tandem duplications to have played the largest role by enriching eight molecular function GO-terms (Figure 6; Table S10). Many of these functions were also enriched in our genome-wide analysis, including acyl-transferase activity, oxidoreductase activity, and endopeptidase inhibitors.

**Figure 6.**
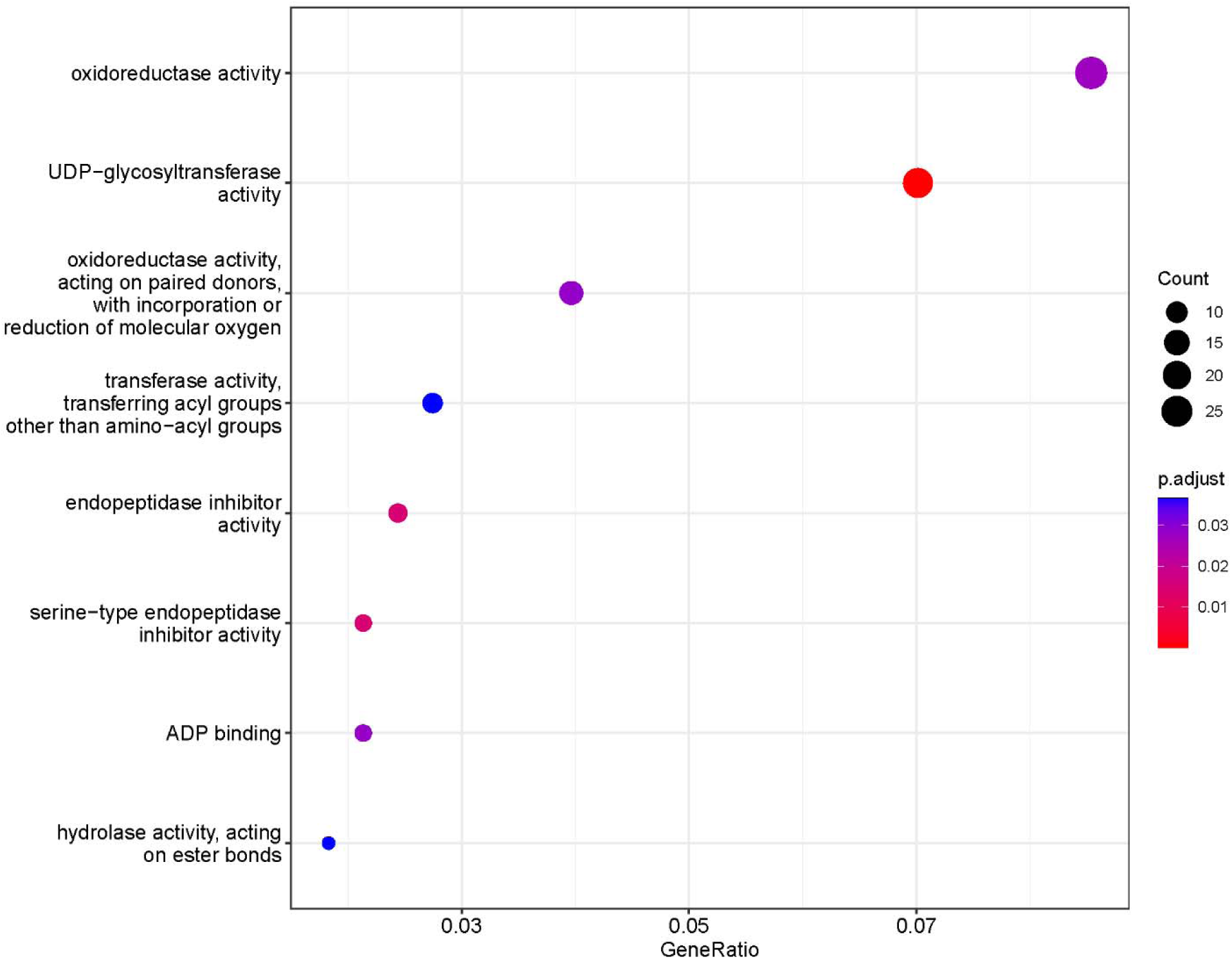
Results of over-representation analysis examining the molecular function GO-terms of differentially expressed genes enriched by tandem duplications. Eight terms were found to be enriched in our dataset, more than any other duplication type (see Table 2 for details).

## Discussion

Our genome assembly produced using only PacBio HiFi reads is highly complete and contiguous. It is, however, important to note that it is not chromosome-scale or completely error-free. Despite great improvements in genome assembly algorithms and long-read sequencing accuracy, additional datasets (e.g. Hi-C) are generally required to produce chromosome-scale gapless assemblies without substantial structural errors [55]. That said, our largest contigs (>100Mb in length), may represent entire chromosomes and were more than sufficient to detect the presence of an ancient whole-genome duplication (Figure 2A). Furthermore, most of the gene content was represented, making our assembly suitable for use in mapping-based analysis such as differential expression studies.

Our differential expression analyses identified thousands of genes that are differentially expressed in *D. wrightii* plants that have been attacked by herbivores, hundreds of which remain highly expressed for multiple days. Many of the DEGs that we identified are known homologs of functionally important genes in other plant-insect systems [54], although it is possible that some of these genes may be induced by mechanical wounding and are not specifically involved in responses to herbivory. The lack of wounding controls in our study design is a limitation, although mechanisms to impose mechanical wounding that accurately mimics the small amount of tissue damaged per “bite,” and accumulated wounding over time remains challenging for most systems (e.g., Mithofer et al. 2005 [58]). Nonetheless, our functional annotation allows us to confidently identify many DEGs as having functional roles in the production of herbivore-induced plant defenses. Furthermore, the presence of genes with long-term (e.g. multiple-day) changes to expression levels is likely indicative of a role in herbivore defense, as prior studies have found that differential expression of wounding-response genes to be most pronounced on short time scales (e.g. less than 24h) [56]. As such, our list of DEGs provides an excellent starting point for future studies that will lead to further insights into the molecular basis of ecological between plants and herbivorous insects over ecologically relevant time periods.

Gene and genome duplications have been shown to play an important role in the arms race between plants and insects [20], but overall, the molecular underpinnings of plant-insect co-evolution remain poorly resolved. By examining the role of various gene duplication types in expanding the functional repertoire of herbivore-induced genes, we have helped to close this gap in our understanding. The finding that tandem duplications are the most important is consistent with previous research that has found biosynthetic genes involved with the production of secondary metabolites to form distinct clusters generated via tandem duplication bursts [57]. Future studies of the duplication dynamics in both plant and insects may begin to unravel the role of biotic selective pressures in generating chemical defense diversity and the resources presented here will serve to accelerate this line of research.

## Concluding Remarks

In sum, we present a high quality long-read genome assembly for the sacred datura plant (*Datura wrightii*). We then analyze its gene duplication history and use it as the basis for a genome-guided analysis of herbivore-induced gene expression changes. Multiple tools supported the presence of a well-documented ancient whole-genome duplication event in this species; our analysis identified thousands of differentially expressed genes, some of which have known functional annotations based on comparison to existing reference genomes. We further show, using a GO-term enrichment approach, that tandem duplications have played an important role in the evolution of *D. wrightii*’s herbivore-responsive gene repertoire. Together, these data provide a valuable resource more broadly and will contribute to future studies of angiosperm evolution and the molecular basis of ecological interactions between plants and herbivorous insects.

## Declarations

### Ethical Approval and consent to participate

No collections were undertaken in protected regions or of protected species, thus no permits were required for fieldwork. In lieu of voucher specimens, observations of the initial plant, alongside the users who identified it, from which seeds were collected in 2016 can be found on iNaturalist (https://www.inaturalist.org/observations/156947070). All methods were carried out in accordance with relevant institutional, national, and international guidelines and legislation.

### Consent to Publication – not applicable

#### Data and Material Availability statement

Assembly and raw HiFi reads can be found on NCBI (BioProject: PRJNA966699; BioSample: SAMN34546691). Supporting datasets, which includes annotation, can be found at https://github.com/caterpillar-coevolution/Datura-wrightii-genome-project alongside scripts used for computational analyses. Raw transcriptome data will be available on NCBI GenBank (BioProject: PRJNA966699). Seeds from the Portal 2016 cohort can be made available on request.

Reviewer link for un-released data: https://dataview.ncbi.nlm.nih.gov/object/PRJNA966699?reviewer=skgd96q1m91kppueafvea7va5p

## Supporting information

Table S1 - Inspector Results

Table S2 - RepeatMasker Results

Table S3 - Annotation Statistics from AGAT

Table S4 - RNAseq read mapping statistics

Table S5 - Pairwise differential expression results

Table S6 - Detailed GSEA results

Table S7 - timecourse expression results

Table S8 - duplication and syntenic depth results

Table S9 - global GO-term enrichment results

Table S10 - Herbivore-induced GO-term results

High resolution version of Figure 4 for improved readability

## Acknowledgments

We thank Professor Luciano Matzkin for his guidance throughout the process of working on the project. We would also like to thank the 2016 staff and volunteers of the Southwestern Research Station for assistance with field work and initial collection of seeds.

## Author contributions

JKG, AO, JDH, and MSB conceived of the project and designed the experiments. JKG collected initial samples for genome assembly. AO and JDH collected samples for differential expression analysis. JKG and MM conducted computational analyses. JKG drafted the initial manuscript. All authors reviewed and contributed to the final version of this manuscript.

