## Supplemental Figures (S1-3) for "A *de novo* long-read genome assembly of the sacred datura plant (*Datura wrightii*) reveals a role of tandem gene duplications in the evolution of herbivore-defense response"

**Supplemental Materials**

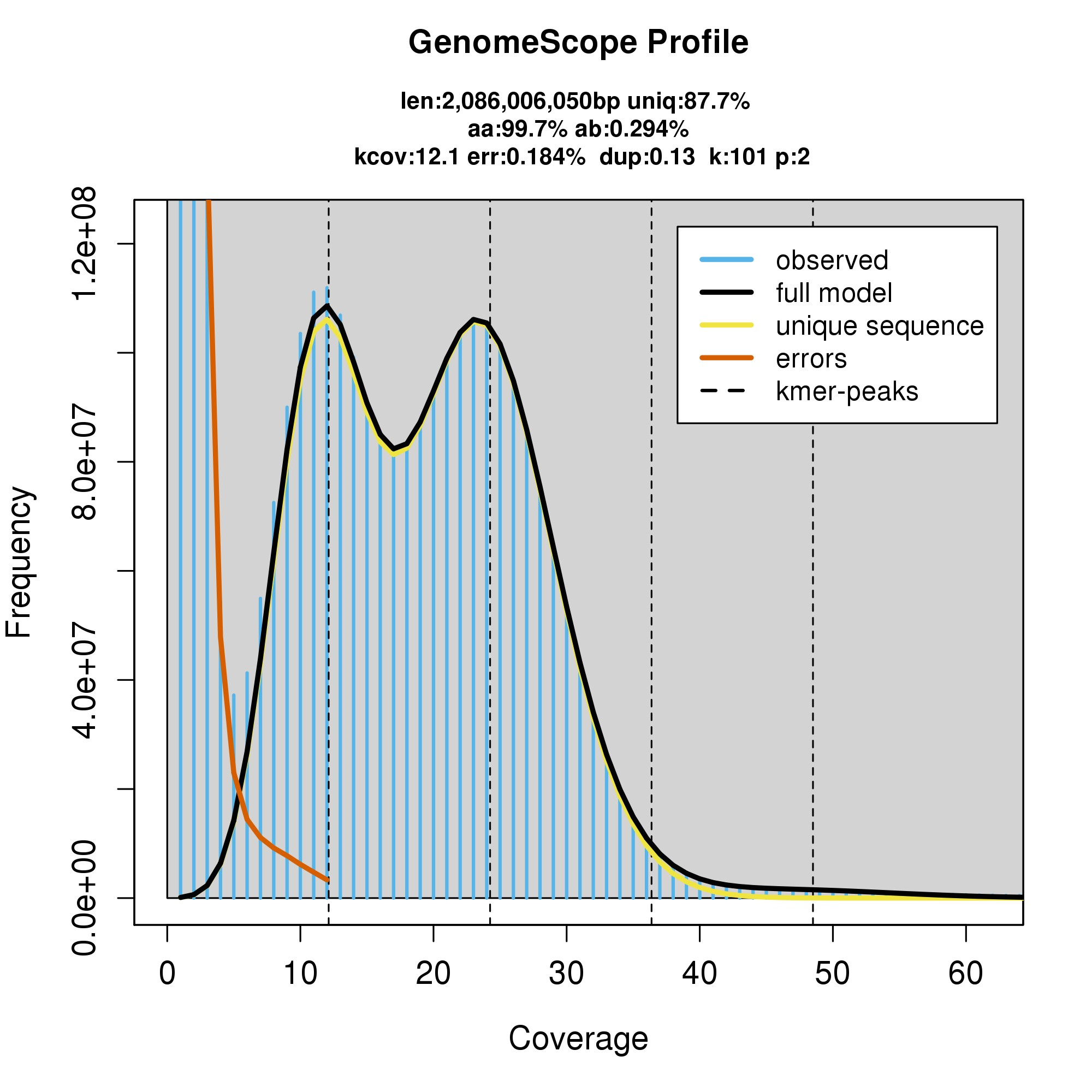

**Figure S1.** Results of kmer analysis to estimate genome size and heterozygosity from raw reads.

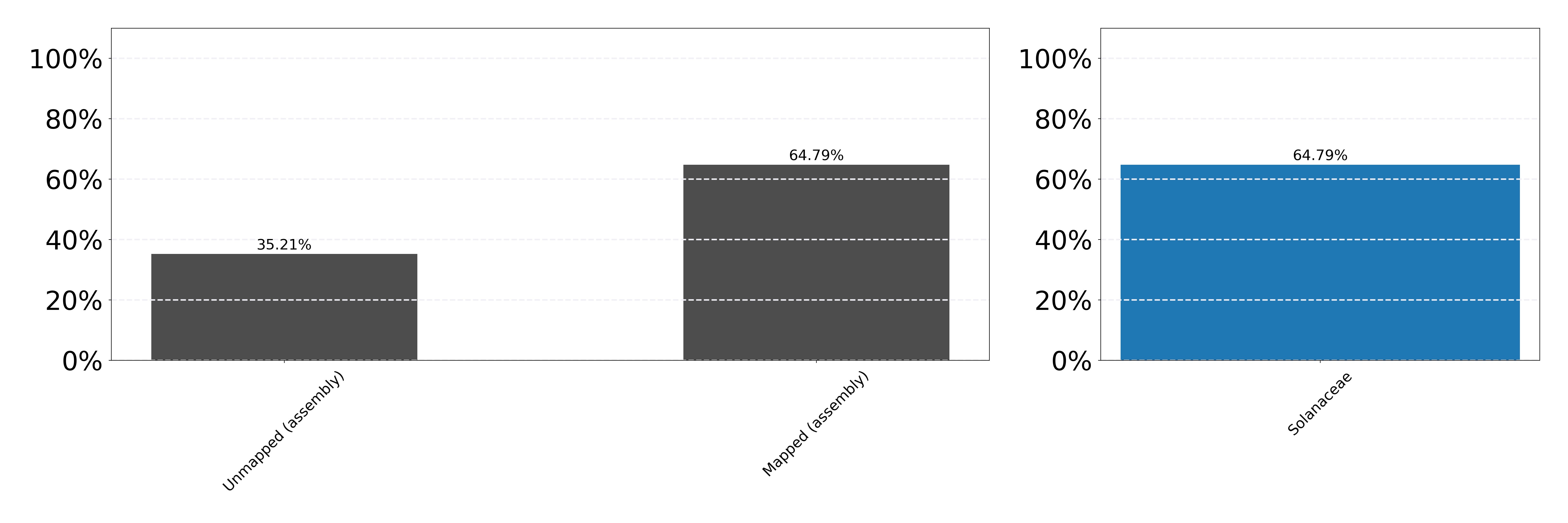

**Figure S2.** Blobtools coverage plot. (left) the proportion of mapped and unmapped reads. (right) the taxanomic identification of mapped reads. All reads were identified as Solanaceae, indicating no contamination in our assembly.

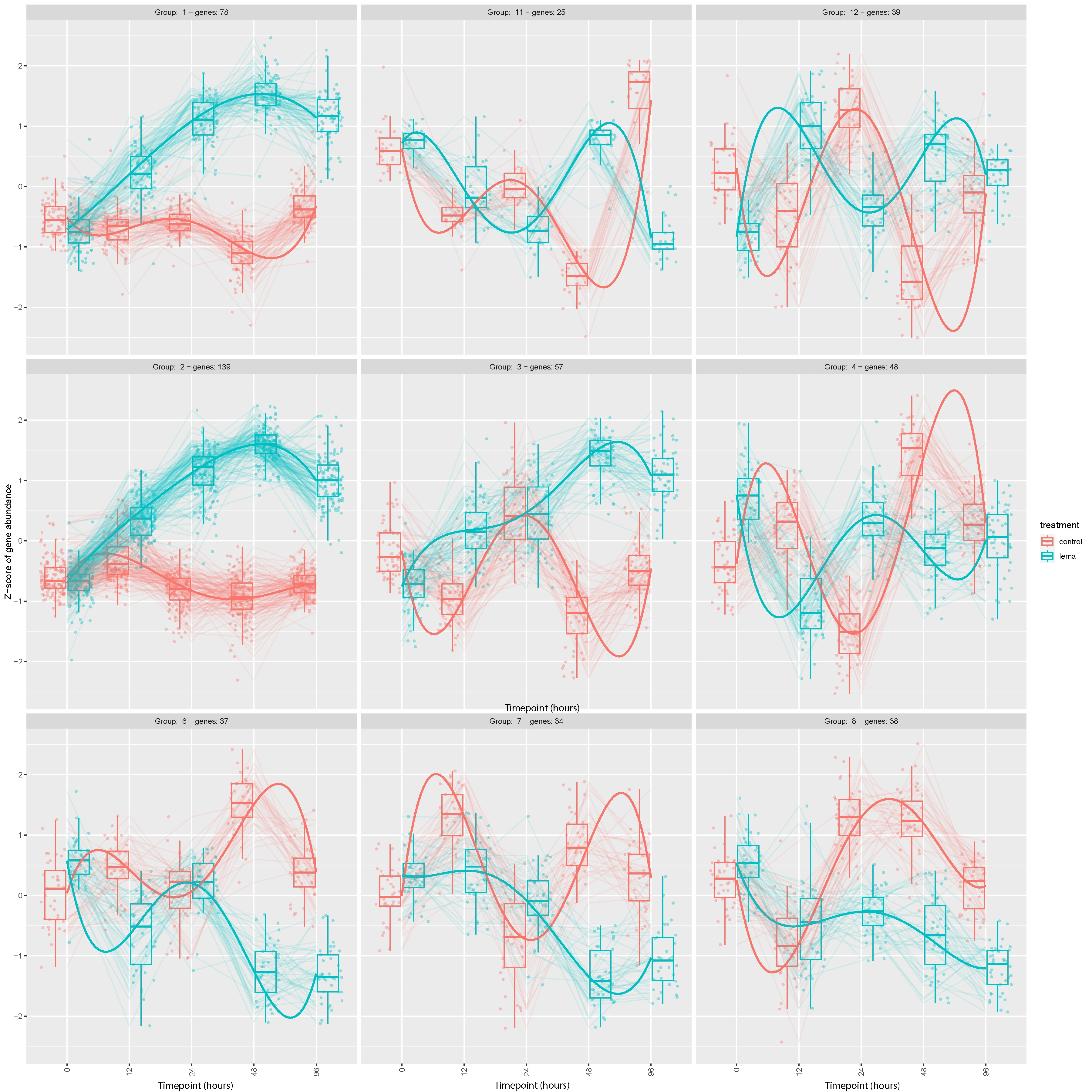

**Figure S3.** Extended results of cluster analysis showing the expression patterns of the nine largest gene clusters identified in our timecourse analysis. Only the two largest gene clusters (also shown in Figure 5) showed clear differences between treatments over time.
