## Supplementary figures and images for "A *de novo* long-read genome assembly of the sacred datura plant (*Datura wrightii*) reveals a role of tandem gene duplications in the evolution of herbivore-defense response"

### High resolution version of Figure 4 for improved readability

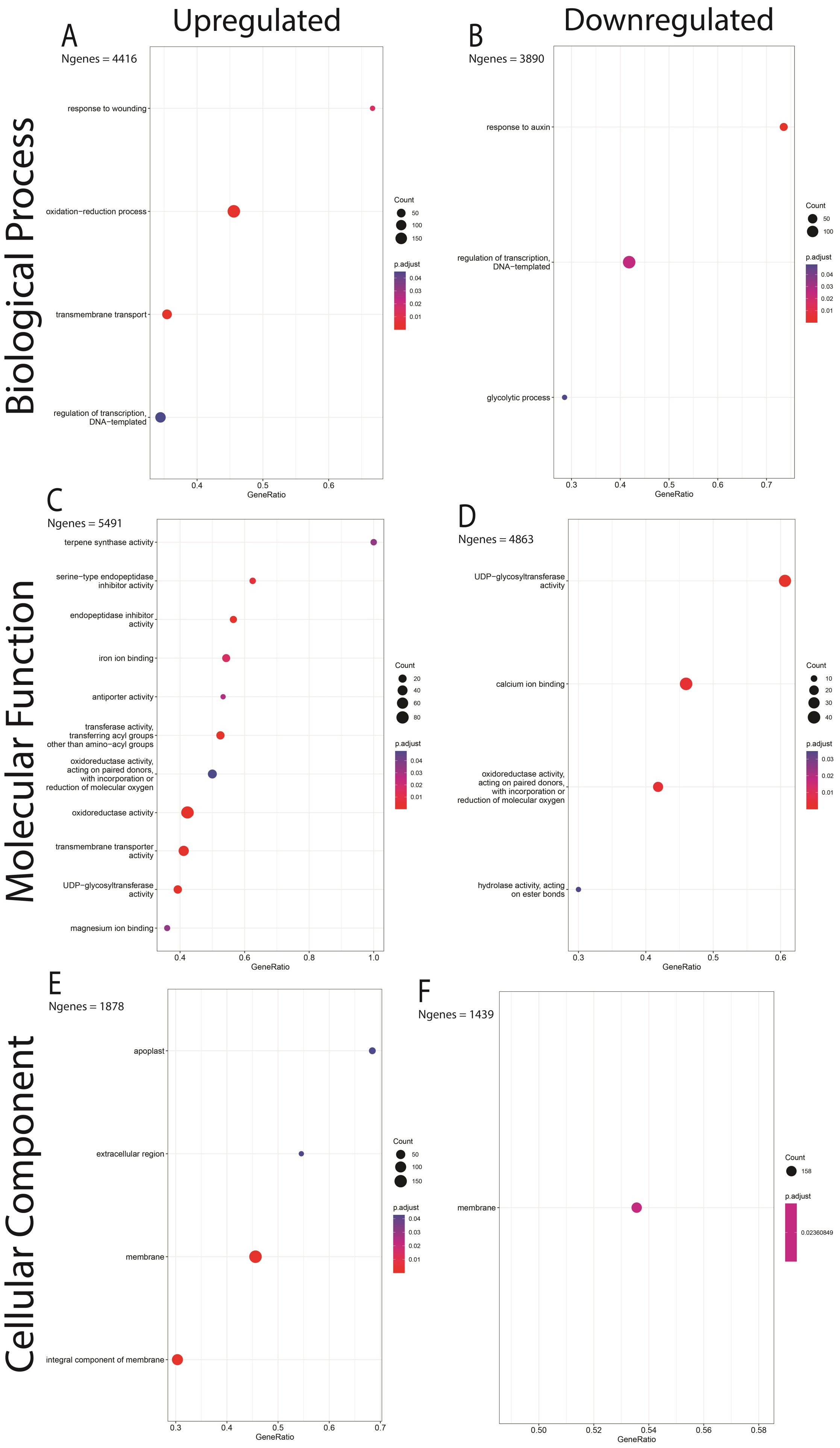
